# Actin-related protein 5 functions as a novel modulator of MyoD and MyoG in skeletal muscle and in rhabdomyosarcoma

**DOI:** 10.1101/2021.05.27.446008

**Authors:** Tsuyoshi Morita, Ken’ichiro Hayashi

**Affiliations:** Department of Biology, Wakayama Medical University, 580 Mikazura, Wakayama, 641-0011, Japan; Department of RNA Biology and Neuroscience, Osaka University Graduate School of Medicine, 2-2 Yamadaoka, Suita, Osaka 565-0871, Japan

## Abstract

Myogenic regulatory factors (MRFs) are pivotal transcription factors in myogenic differentiation. MyoD commits cells to the skeletal muscle lineage by inducing myogenic genes through recruitment of chromatin remodelers to its target loci. This study showed that Actin-related protein 5 (Arp5) acts as an inhibitory regulator of MyoD and MyoG by binding to their cysteine-rich (CR) region, which overlaps with the region essential for their epigenetic functions. Arp5 expression was faint in skeletal muscle tissues. Excessive Arp5 in mouse hind limbs caused skeletal muscle fiber atrophy. Further, Arp5 overexpression in myoblasts inhibited myotube formation by diminishing myogenic gene expression, whereas Arp5 depletion augmented myogenic gene expression. Arp5 disturbed MyoD-mediated chromatin remodeling through competition with the three-amino-acid-loop-extension-class homeodomain transcription factors the Pbx1–Meis1 heterodimer for binding to the CR region. This antimyogenic function was independent of the INO80 chromatin remodeling complex, although Arp5 is an important component of that. In rhabdomyosarcoma (RMS) cells, Arp5 expression was significantly higher than in normal myoblasts and skeletal muscle tissue, probably contributing to MyoD and MyoG activity dysregulation. Arp5 depletion in RMS partially restored myogenic properties while inhibiting tumorigenic properties. Thus, Arp5 is a novel modulator of MRFs in skeletal muscle differentiation.

## Introduction

Actin-related protein 5 (Arp5), encoded by *Actr5*, is a nuclear-localized actin-like protein (Schafer and Schroer, 1999). Studies have investigated the role of Arp5 in the nucleus as one of the subunits of the ATPase-dependent chromatin remodeling complex INO80 (Shen et al., 2000). INO80 regulates various DNA metabolic processes, such as gene expression, DNA replication, and DNA repair by nucleosome sliding (Polil et al., 2017). It contains β-actin and three Arp family members (Arp4, Arp5, and Arp8). Arp5 forms the Arp5 module with Ies6 (encoded by *Ino80c*), which is necessary for ATP hydrolysis and nucleosome sliding by INO80 (Yao et al., 2015). However, a few features of Arp5 are unrelated to INO80. In *Arabidopsis*, Arp5 and Ino80, an essential ATPase component of INO80, have common and distinct features in plant growth and development (Kang et al., 2019). In addition, Arp5 plays an INO80-independent role in regulating the differentiation of vascular smooth muscle cells (SMCs) (Morita et al., 2014). In rat SMCs, Arp5 expression is strongly inhibited, although INO80 activity is generally necessary for cell growth and proliferation. Arp5 inhibits SMC differentiation by interacting with and inhibiting the SAP family transcription factor myocardin, which is a key regulator for the induction of SMC-specific contractile genes, indicating that low Arp5 expression in SMCs contributes to maintaining their differentiation status. In contrast, in cardiac and skeletal muscles, although Apr5 expression is low, its physiological significance is unclear.

Myogenic regulatory factors (MRFs), such as MYF5, MyoD, MyoG, and MRF4, are skeletal-muscle-specific basic helix–loop–helix (bHLH) transcription factors and master regulators of skeletal muscle development. They recognize a *cis*-regulatory element E-box, usually found in promoter and enhancer regions of muscle-specific genes, as a heterodimer with ubiquitous bHLH proteins of the E2A family (E12 and E47) (Funk et al., 1991). MRFs enhance the transcriptional activity of myogenic genes via chromatin remodeling by recruiting the switch/sucrose nonfermentable (SWI/SNF) complex to previously silent target loci (Roy et al., 2002; Ohkawa et al., 2007). This epigenetic activity depends on a histidine- and cysteine-rich (H/C) region N-terminal to the bHLH domain in MRFs (Gerber et al., 1997). This region contains a CL-X-W motif, which is a binding site for the heterodimer of three-amino-acid-loop-extension (TALE)-class homeodomain transcription factors (Pbx1 and Meis/Prep1) (Knoepfler et al., 1999). Funk and Wright (1992) reported that E-box elements recognized by MyoG are occasionally flanked by a novel consensus motif TGATTGAC, which was also identified as a binding motif for the Pbx1–Meis/Prep1 heterodimer (Knoepfler et al., 1999). Thus, MRFs form a complex with the TALE heterodimer on DNA, leading to the recruitment of chromatin remodelers and an increase in the accessibility of their target loci.

This study reported a novel role of Arp5 in myogenic differentiation of skeletal muscle cells. We identified Arp5 as an inhibitory binding protein for MyoD and MyoG in skeletal muscle and rhabdomyosarcoma (RMS) cells. Results showed that Arp5 competes with the Pbx1–Meis1 heterodimer for interaction with the cysteine-rich (CR) region of MyoD and MyoG and consequently inhibits myogenic differentiation.

## Results

### Arp5 prevents skeletal muscle development by inhibiting MRF expression

Arp5 expression was significantly low in heart, aorta, and especially, hind limb muscle tissues (Figure 1A). Besides, Arp5 expression in primary mouse myoblasts significantly decreased with myotube differentiation (Figure 1B). The public human transcriptome databases (Human Protein Atlas [HPA], Genotype-Tissue Expression [GTEx], and Functional Annotation of the Mouse/Mammalian Genome 5 [FANTOM5]) showed low *ARP5* expression in skeletal muscle tissues (Figure 1—figure supplement 1). In this study, 5 weeks after injection of the Arp5-AAV6 vector, the muscle fiber thickness significantly reduced compared to the control group (Figure 1C). In these atrophic muscles, gene expression levels of MRFs (*Myod1, MyoG*, and *Myf6*) significantly decreased, accompanied by a decrease in other skeletal muscle markers, such as *Myh4, Acta1*, and *Tnni1* (Figure 1D).

**Figure 1.**
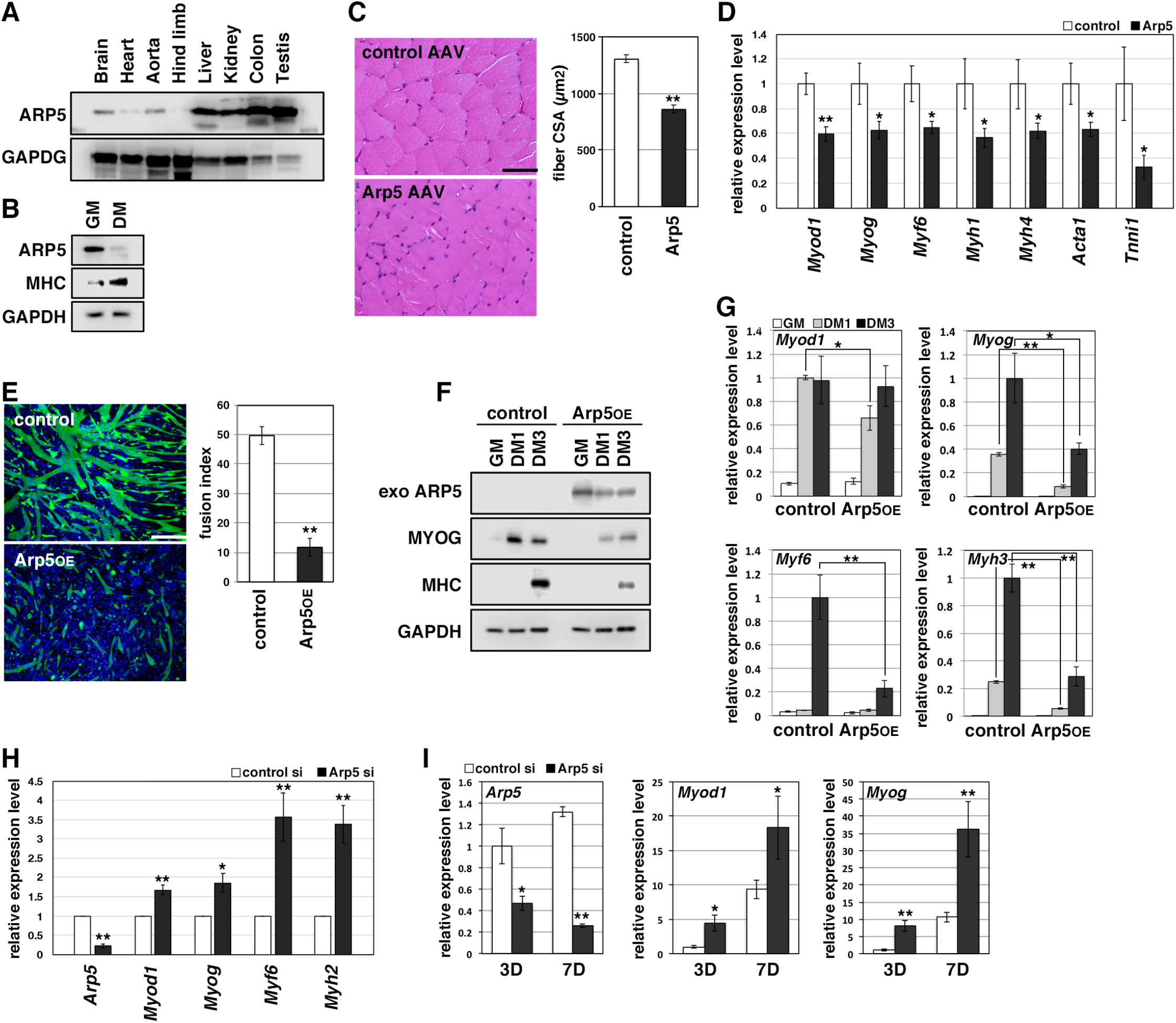
Actin-related protein 5 (Arp5) inhibits skeletal muscle differentiation. (A) Arp5 expression in mouse tissues. (B) Arp5 and myosin heavy chain (MHC) expression in C2C12 cells cultured in growth medium (GM) or differentiation medium (DM). (C) Representative images of hematoxylin and eosin (H&E)-stained section of the hind limb muscle from mice injected with control or Arp5-AAV6 vector (left). Scale bar = 50 µm. Muscle fiber cross-sectional area (CSA) measured in 190 fibers and statistically analyzed (right). (D) Myogenic gene expression in AAV6-vector-injected hind limb muscles. (E) Representative fluorescence images of differentiated C2C12 cells transfected with green fluorescent protein (GFP) alone (control) or together with Arp5 (Arp5^OE^) (left). Nuclei were visualized by Hoechst 33342. Scale bar = 100 µm. The fusion index was measured on 22 images and statistically analyzed (right). (F) Myogenic protein expression in C2C12 cells transfected with control or Arp5 expression vector. The cells were cultured in GM or DM for 1 day (DMl) or 3 days (DM3) after transfection. (G) Myogenic gene expression in Arp5-transfected C2C12 cells. (H) Myogenic gene expression in mouse primary myoblasts transfected with control or Arp5 short interfering RNA (siRNA). (I) Myogenic gene expression in l0Tl/2 cells treated with 5-azacytidine. The cells were transfected with control or Arp5 siRNA prior to 5-azacytidine treatment. All statistical data are presented as the mean ± standard error of the mean (SEM). *\*P* < 0.05, *\*\*P* < 0.01 (Student’s t-test).

In C2C12 cells, Arp5 overexpression significantly inhibited the fusion ability of myoblasts and the induction of MyoG and myosin heavy chain (MHC) under the conditions for myotube formation (Figure 1E,F). In addition, the induction of *Myod1, MyoG, Myf6*, and *Myh3* was also significantly inhibited (Figure 1G). Conversely, gene silencing of Arp5 by short interfering RNA (siRNA) (Arp5-si) increased MRF and *Myh2* expression (Figure 1H).

Long-term exposure to 5-azacytidine converts C3H 10T1/2 mouse embryo fibroblasts into differentiated skeletal muscle cells with the induction of endogenous MRF expression (Davis et al., 1987). There was significant induction of endogenous *Myod1* and *MyoG* 7 days after 5-azacytidine treatment, and Arp5 silencing significantly augmented *Myod1* and *MyoG* induction (Figure 1I). These findings show that Arp5 plays an inhibitory role in skeletal muscle development via regulating MRF expression and activity.

### High Arp5 expression contributes to defective myogenic differentiation in rhabdomyosarcoma

RMS is a common soft-tissue sarcoma developed from skeletal muscle with defective myogenic differentiation. High MyoD and MyoG expression is used in the clinical diagnosis of RMS, but they are not fully active for the induction of their target genes (Folpe 2002; Keller and Guttridge 2013). *ARP5* expression was significantly higher in human embryonal and alveolar RMS tissues compared with healthy and tumor-adjacent skeletal muscles (Figure 2A) and in RD cells compared with primary human myoblasts (Figure 2B), indicating that Arp5 overexpression contributes to MRF dysfunction in RMS.

**Figure 2.**
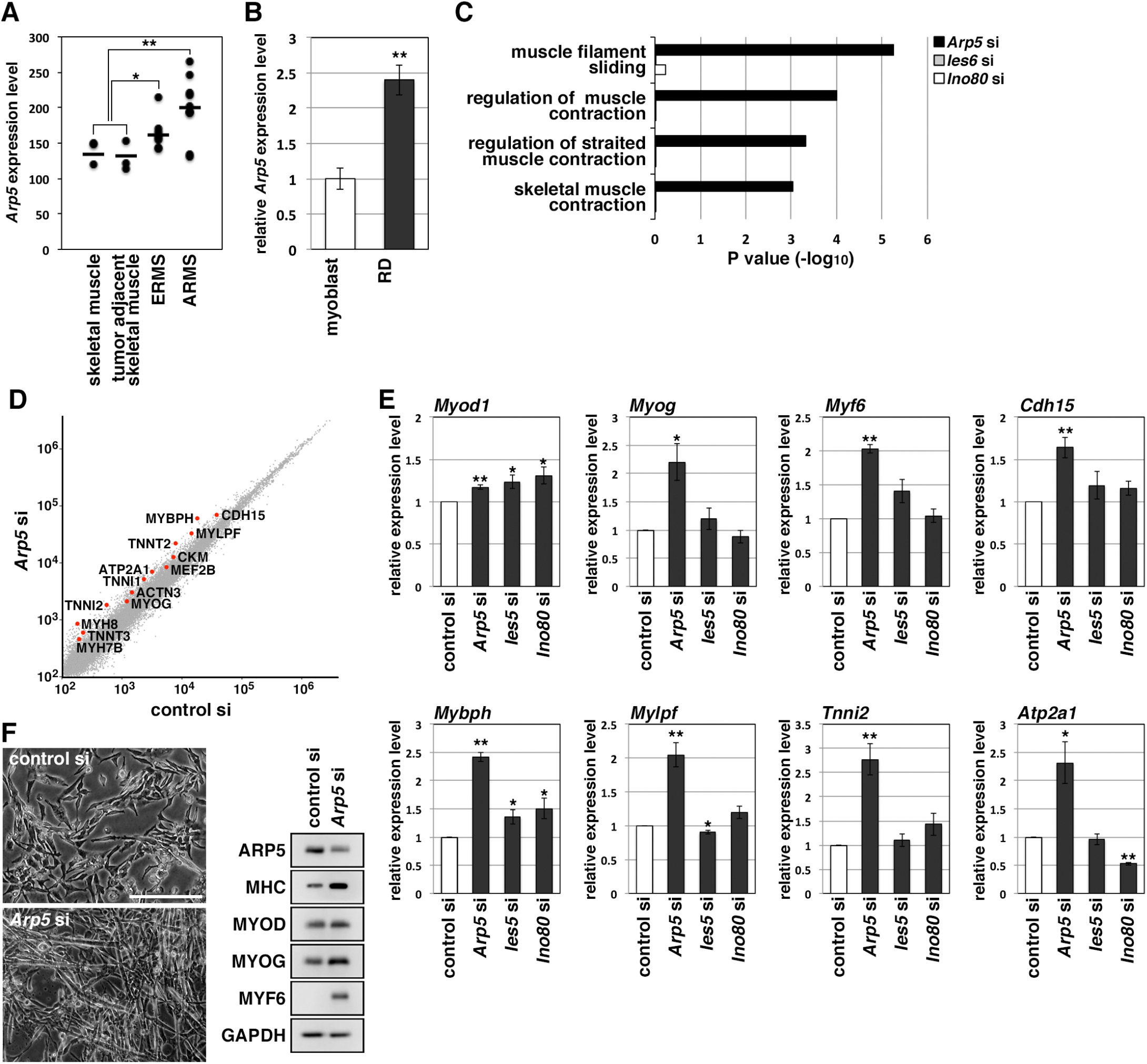
Actin-related protein 5 (Arp5) knockdown increases myogenic gene expression in rhabdomyosarcoma (RMS) cells. (A) *Arp5* expression in normal skeletal muscle, tumor-adjacent skeletal muscle, embryonal RMS (ERMS), and aveolar RMS (ARMS). Bars indicate average expression levels. (B) *ARP5* expression in human primary myoblasts and RD cells. (C) Enrichment analysis of muscle-related Gene Ontology terms from DNA microarray data on comparison between control-si and Arp5-, Ies6-, or Ino80-si samples. (D) Scatter plot of gene expression level in control-si and Arp5-si RD cells. (E) Myogenic gene expression in control-, Arp5-, Ies6-, and Ino80-si RD cells. (F) Representative phase-contrast images of control-si and Arp5-si RD cells (left). Scale bar = 100 µm. Myogenic protein expression in these cells (right). All statistical data are presented as the mean ± standard error of the mean (SEM). \**P* < 0.05, *\*\*P* < 0.01 (Student’s t-test).

Microarray analysis on Arp5-silencing RD cells showed that Arp5-si increased the expression of numerous genes involved in muscle function and development (Figure 2C,D, Figure 2—Table supplement 1). The increased myogenic gene expression was also confirmed by real-time reverse transcription polymerase chain reaction (RT-PCR) (Figure 2E). Unlike myoblasts, RD cells have little or no ability to form myotubes, even under serum-free culture conditions, but Arp5 depletion led RD cells to form numerous myotube-like structures with upregulation of myogenic marker proteins (Figure 2F).

Arp5 is one of the subunits of INO80, so siRNA (Ies6-si and Ino80-si) also depleted the expression of other subunits (Ies6 and Ino80). The genes altered by the silencing of each of these partly overlapped (Figure2—figure supplement 1A,B); 39.2% and 32.4% of the genes altered by Arp5-si overlapped with those altered by Ies6-si and Ino80-si, respectively. Contrary to Arp5-si, however, Ies6-si and Ino80-si hardly increased the expression of muscle-related genes (Figure 2C,E, Figure 2—figure supplement 1C–E). These findings show that Arp5 overexpression inhibits the expression of muscle-related genes and, therefore, terminal myogenic differentiation in RMS in an INO80-independent manner.

### Arp5 knockout leads to loss of tumorigenicity of RD cells

In Arp5-knockout RD (Arp5-KO) cells, a few nucleotides downstream of the start codon of *Arp5* were deleted by CRISPR-Cas9 genome editing, causing Arp5 deletion (Figure 3A). Arp5-KO cells showed significant upregulation of many kinds of skeletal muscle-related genes (Figure 3B).

**Figure 3.**
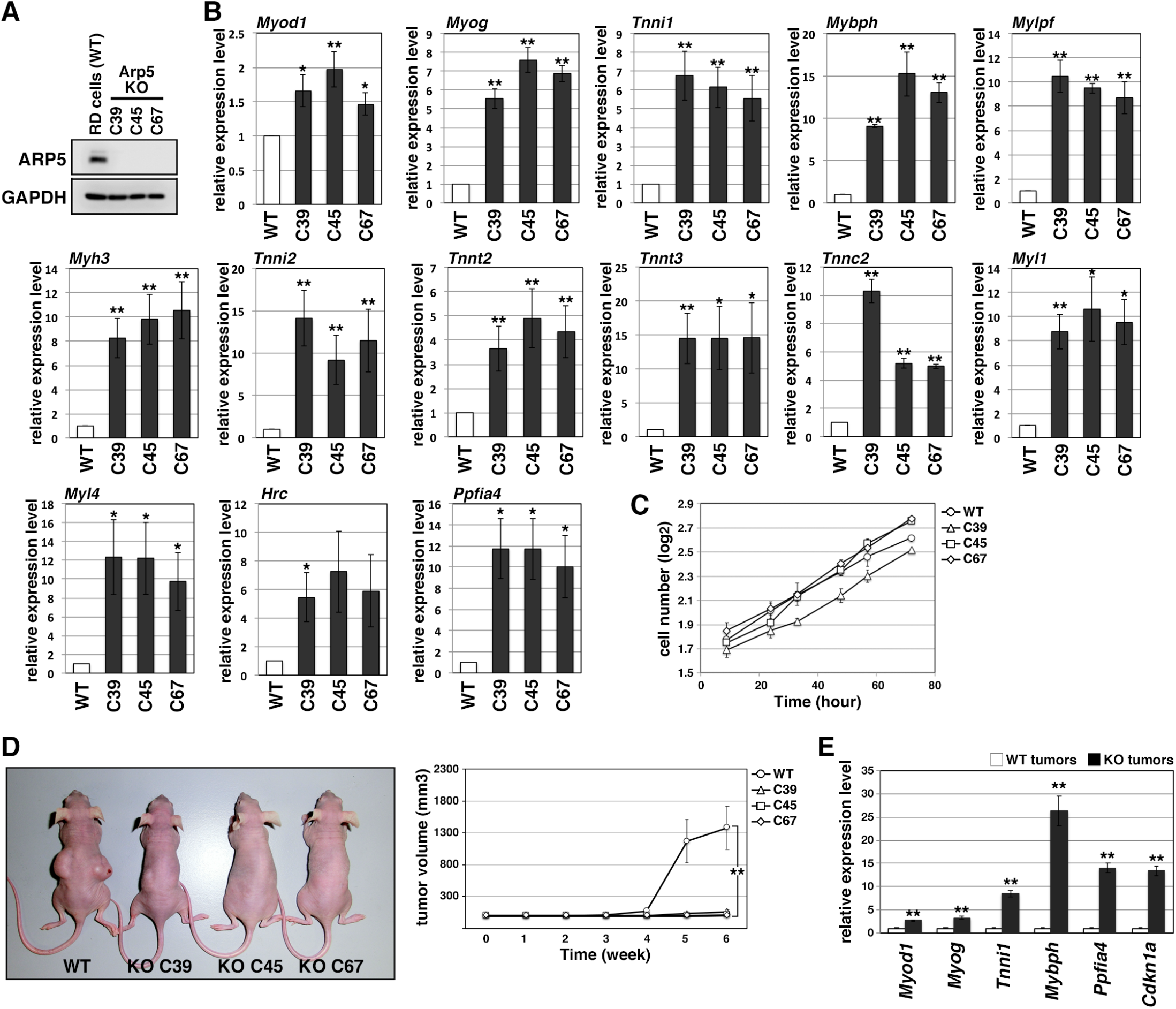
Actin-related protein 5-knockout (Arp5-KO) RD clones show increased expression of myogenic genes and decreased tumorigenicity. (A) Arp5 expression in three individual clones of Arp5-KO cells (C39, C45, and C67) and their parental RD cells (wild-type [WT]). (B) Myogenic gene expression in WT and Arp5-KO cells. (C) Growth curve of WT and Arp5-KO cells. (D) Xenograft model of WT and Arp5-KO cells in nude mice (left). Tumor volumes measured every week after inoculation and statistically analyzed (right). (E) Myogenic gene expression in xenograft tumors. All statistical data are presented as the mean± standard error of the mean (SEM). *\*P* < 0.05, *\*\*P* < 0.01 (Student’s t-test).

Under culture conditions, Arp5-KO cells proliferated as well as parental RD cells (doubling time of wild-type [WT] RD cells [*t*_2_(WT)] = 29.7 h, *t*_2_(C39) = 29.8 h, *t*_2_(C45) = 23.8 h, *t*_2_(C67) = 26.7 h; Figure 3C). However, when transplanted subcutaneously into nude mice, they rarely formed tumor nodules differently from parental cells (Figure 3D). These rare tumor nodules showed higher expression of muscle-related genes and *Cdkn1a*, which encodes a cyclin-dependent kinase inhibitor p21^WAF1/CIP1^ controlling cell cycle arrest and myogenic differentiation during muscle development and regeneration (Halevy et al., 1995; Figure 3E). Thus, Arp5 deletion partially restores the impaired myogenic differentiation potential of RD cells and inhibits their tumorigenesis in vivo.

### Arp5 inhibits MyoD and MyoG activity through binding to their cysteine-rich region

MRF transcription is regulated by positive autoregulation (Tapscott SJ, 2005). To examine whether Arp5 inhibits the activity or transcription of MRF, immunoprecipitation and reporter promoter analysis were performed. Immunoprecipitation showed that Arp5 binds to both MyoD and MyoG but not to the ubiquitous bHLH protein E47 (Figure 4A). MyoG was precipitated more efficiently than MyoD with Arp5. The amino acid sequences of basic and HLH regions were highly conserved between MyoD and MyoG (Figure 4—figure supplement 1), but these domains were not necessary for their interaction (Figure 4B). The CR region, a part of the H/C region, was also highly conserved between them (Figure 4—figure supplement 1), and CR region deletion mostly abolished the Arp5-binding ability of both MyoD and MyoG (Figure 4B,C). MyoG promoter-controlled luciferase reporter assay demonstrated that MyoD strongly enhances MyoG promoter activity, while Arp5 significantly inhibits it (Figure 4D). MyoD ΔCR, in which the CR region was deleted, also increased MyoG promoter activity but to a lesser extent, and this activation was completely unaffected by Arp5 (Figure 4D), indicating that the direct interaction with MyoD via the CR region is necessary for Arp5 to inhibit MyoD activity.

**Figure 4.**
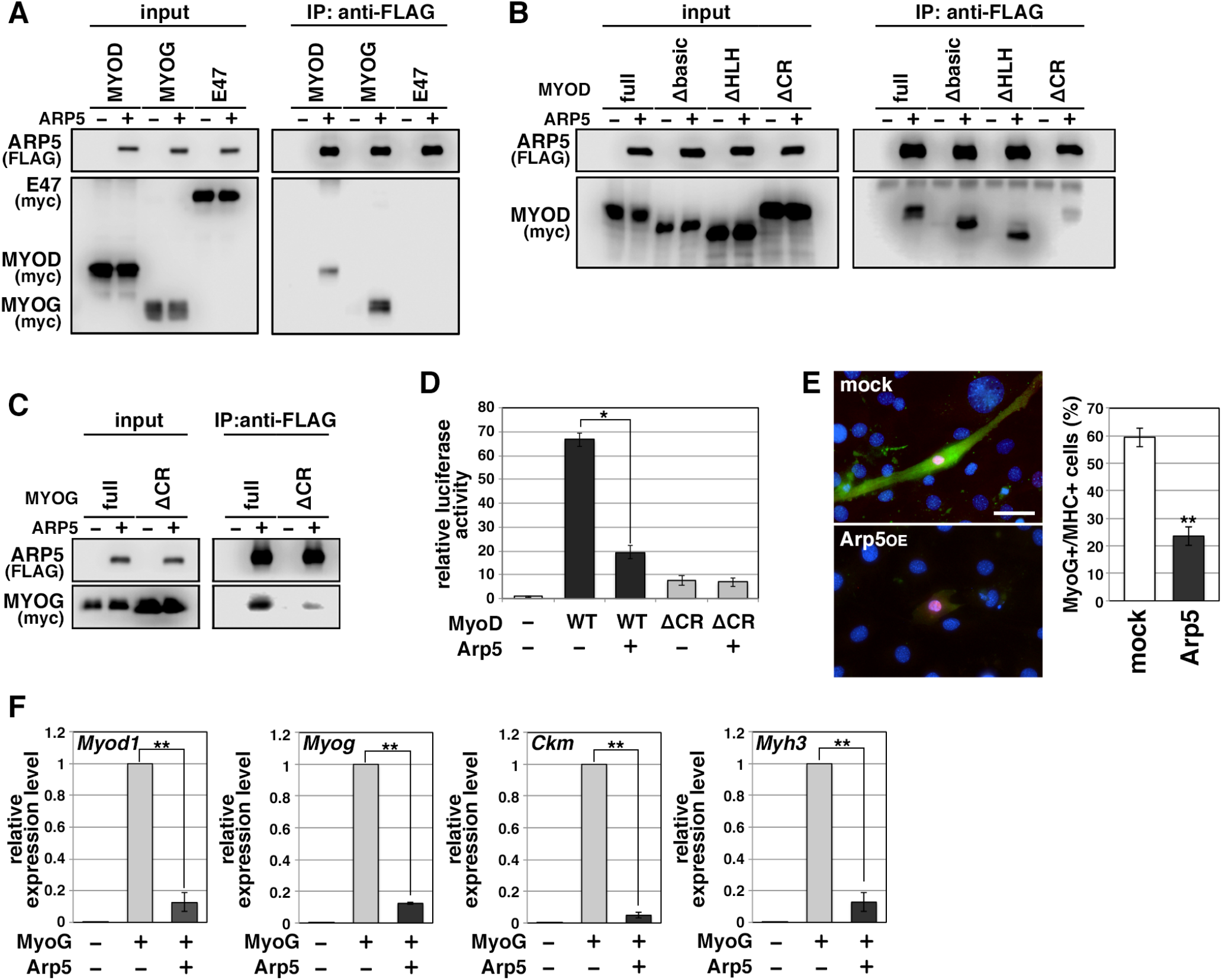
Actin-related protein 5 (Arp5) inhibits the activity of MyoD and MyoG through direct interaction. (A) Co-immunoprecipitation assay between Arp5 and basic helix loop-helix (bHLH) transcription factors MyoD, MyoG, and E47. (B) Co-immunoprecipitation assay between Arp5 and truncated series of MyoD. (C) Co-immunoprecipitation assay between Arp5 and the CR-region-deleted MyoG. (D) MyoG promoter-controlled luciferase reporter assay in C2C12 cells. (E) Representative fluorescence images of l0T/2 cells transfected with MyoG and Arp5 (left). The cells were immunostained with anti-MyoG (red) and anti-myosin heavy chain (MHC, green) antibodies. Nuclei were visualized by Hoechst 33342 (blue). Scale bar = 50 µm. The percentage of MyoG+ MHC+ double-positive cells in MyoG+-positive cells was calculated in 786 cells and statistically analyzed (right). (F) Endogenous myogenic gene expression in l0Tl/2 cells transfected with MyoG and Arp5. All statistical data are presented as the mean± standard error of the mean (SEM). *P < 0.05, **P < 0.01 (Student’s t-test).

Ectopic MyoG expression differentiated 10T1/2 cells into MHC-positive myogenic cells with induction of endogenous myogenic marker genes such as *Myod1, MyoG, Ckm*, and *Myh3* (Figure 4E,F). Co-expression of Arp5 with MyoG, however, significantly decreased the frequency of MHC-positive cells and significantly inhibited the induction of myogenic genes (Figure 4E,F). Thus, Arp5 inhibits MyoD/MyoG activity through direct interaction via their CR region.

### Arp5 competes with Pbx1–Meis1 for binding to the cysteine-rich region of MyoD/MyoG

The H/C region of MyoD is an essential region for its chromatin remodeling activity and effective induction of target genes (Gerber et al., 1997). Pbx1 and Meis1 homeobox proteins are promising candidates as mediators between MyoD and chromatin remodelers because MyoD binds to Pbx1 and Meis1 via the H/C region (Knoepfler et al., 1999). MyoD and the Pbx1–Meis1 heterodimer interacted in the absence of the intermediary DNA; furthermore, this interaction was interrupted by Arp5 (Figure 5A).

**Figure 5.**
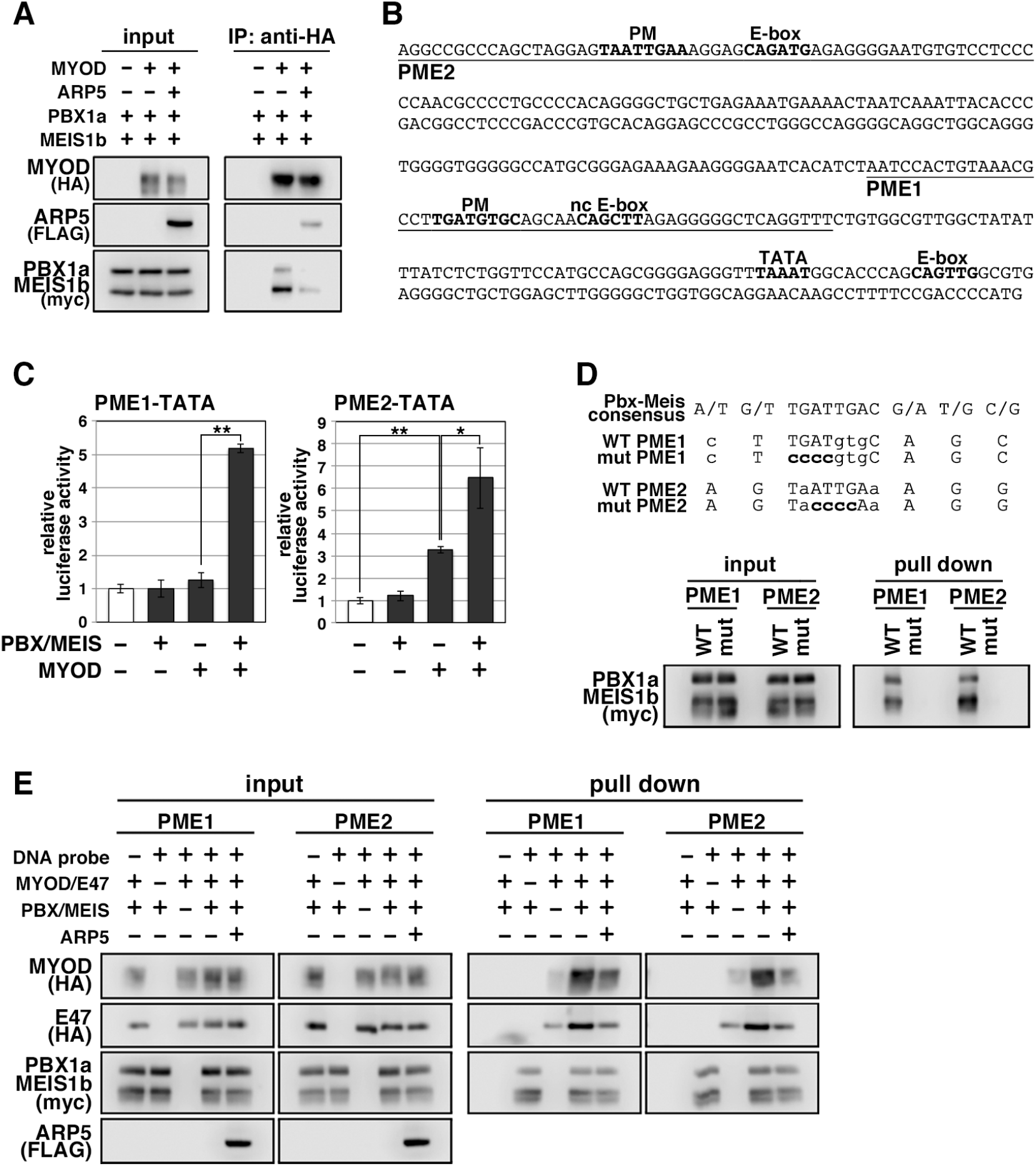
Actin-related protein 5 (Arp5) disturbs the interaction between MyoD and the Pbxl-Meisl heterodimer. (A) Co-immunoprecipitation assay between MyoD and Pbxla Meislb. Co-incubation with Arp5 protein (lanes 3 and 6) diminished their interaction. (B) Sequence of the proximal promoter region of human MYOG. The core sequences of the indicated cis-regulatory elements are highlighted in bold. Pbxl Meisl heterodimer-binding motif/noncanonical E-box (PM/ncE)-containing regions (PMEl and PME2) are underlined. (C) PMEl/2-TATA-controlled luciferase reporter assay in C2C12 cells. (D) Pull-down assay of PME1/2 DNA probes with Pbx1a Meis1b protein. The consensus sequence ofthe PM motif is presented. The mutated nucleotides in the mut PMEl/2 probes are highlighted in bold. (E) Pull-down assay of PMEl/2 DNA probes with MyoD, E47, Pbxla Meislb, and Arp5 proteins.

The *MyoG* locus has been most analyzed for MyoD-mediated chromatin remodeling. The proximal promoter region of *MyoG* reportedly contains two flanking sequences, Pbx1–Meis1 heterodimer-binding motif (PM) and noncanonical E-box (ncE) (Berkes et al., 2004; Figure 5B). When this regulatory region (PM/ncE-containing region 1 [PME1]) was fused to the TATA minimal promoter, the PME1–TATA construct was activated by the combination of Pbx1, Meis1, and MyoD (Figure 5C, left). Upstream of PME1, a novel predicted regulatory region containing PM and E-box motifs (PME2) was identified (Figure 5B). A pull-down assay using DNA-probe-conjugated beads of the PMEs demonstrated that the Pbx1–Meis1 heterodimer recognizes the PM motif of both PME1 and PME2 and that this interaction completely disappears by disruption of the PM motif via mutagenesis (Figure 5D). The PME2–TATA construct was also synergically activated by MyoD and the Pbx1–Meis1 heterodimer (Figure 5C, right). Thus, both PME1 and PME2 are functional *cis*-regulatory elements for MyoD and the Pbx1–Meis1 heterodimer.

The DNA–protein pull-down assay also demonstrated that the MyoD–E47 heterodimer weakly binds to PME1 and PME2 DNA probes, which is significantly augmented by co-incubation with the Pbx1–Meis1 heterodimer (Figure 5E). The Pbx1– Meis1 heterodimer was constantly associated with the PMEs regardless of whether MyoD-E47 was present. Arp5 interrupted the augmented interaction of MyoD–E47 with the PMEs but did not affect the interaction of the Pbx1–Meis1 heterodimer with the PMEs (Figure 5E).

We also identified the predicted PME element in the proximal promoter region of *Myf6*, which contains Pbx- and Meis-binding motifs separated by six nucleotides (acTGATgctccaTGACag) close to noncanonical E-boxes (Figure 6A). Jacobs et al. (1999) reported that this type of gapped PM site is also functional for the Pbx1– Meis1-binding site, and we observed significant interaction between the Pbx1–Meis1 heterodimer and *Myf6*’s PM motif (Figure 6B). Similar to PMEs in the *MyoG* promoter, MyoG bound to *Myf6*’s PME synergically with the Pbx1–Meis1 heterodimer, while Arp5 interrupted this binding (Figure 6C). These findings show that Arp5 attenuates MyoD/MyoG recruitment to PME-containing myogenic enhancer regions by disturbing the interaction between MyoD/MyoG and the Pbx1–Meis1 heterodimer.

**Figure 6.**
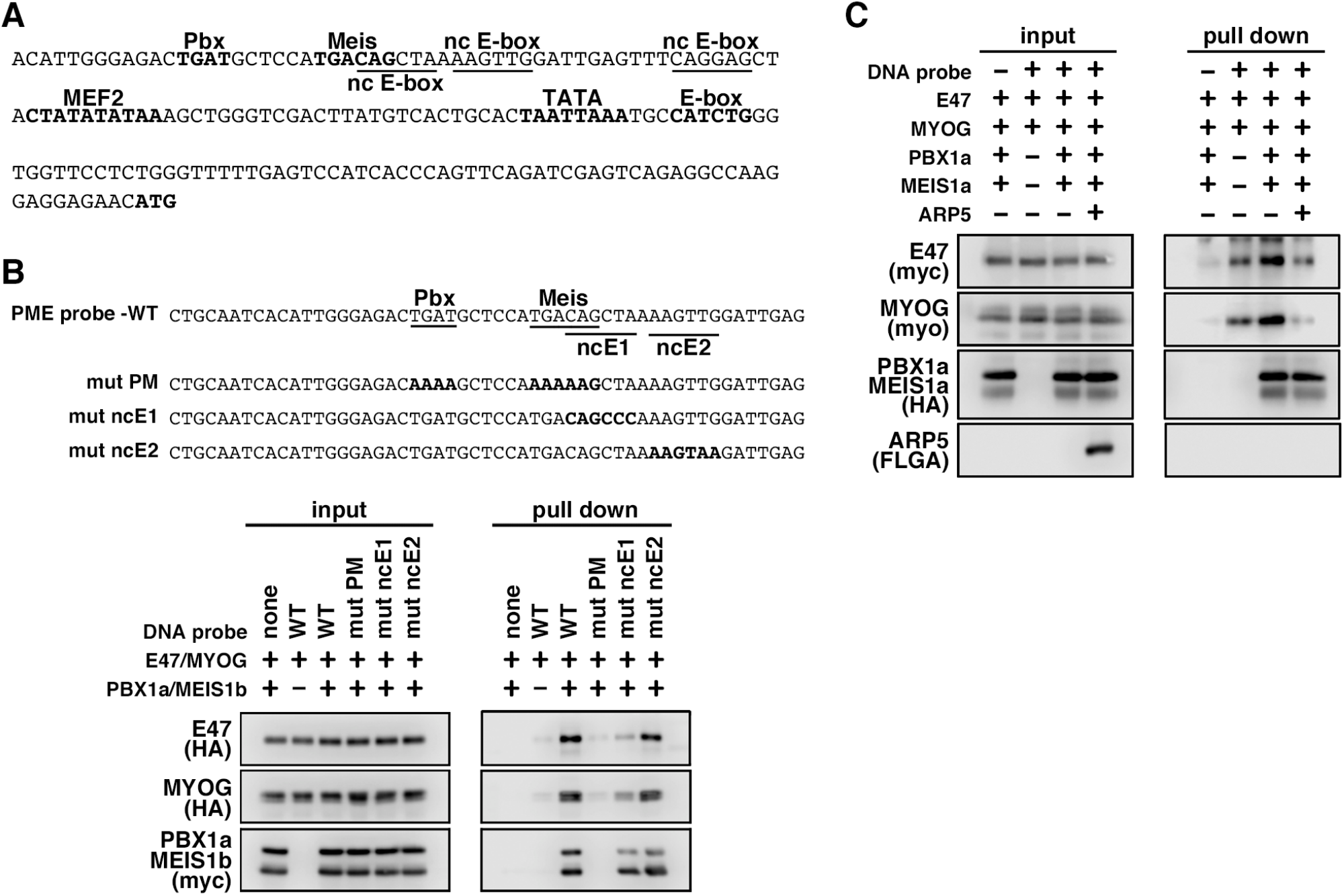
MyoG and the Pbxl-Meisl heterodimer recognize the proximal promoter region of human myogenic regulatory factor 6 (MRF6). (A) Sequence of the proximal promoter region of human *MRF6*. The core sequences of the indicated cis-regulatory elements are highlighted in bold. The core sequences of putative noncanonical E-box motif (nc E-box) are underlined. (B) Pull-down assay of Mrf6 Pbxl-Meisl heterodimer-binding motif/noncanonical E-box (PM/ncE)-containing region (PME) probes with MyoG, E47, and Pbxla Meislb proteins. The mutated nucleotides in the mut PME probes (mut PM, mut ncEl, and mut ncE2) are highlighted in bold (top). The Pbxl Meisl heterodimer recognized the gapped Pbx Meis-binding motif, while MyoG E47 bound to the ncEl site with the Pbx Meis complex (bottom). (C) Pull-down assay of Mrf6 PME probes with MyoG, E47, Pbxla Meislb, and Arp5 proteins.

### Arp5 prevents Brg1–SWI/SNF complex recruitment to MyoD/MyoG target loci

During MyoD-mediated chromatin remodeling, the Pbx1–Meis1 heterodimer is believed to be involved in the recruitment of Brg1-based SWI/SNF chromatin remodeling complex to MyoD target loci as a pioneering factor (de la Serna et al., 2001). Serna et al. (2005) reported that the induction of approximately one-third of MyoD target genes depends on the Brg1–SWI/SNF complex. Therefore, we investigated the change in MyoD target gene expression by Arp5-si in terms of Brg1 dependency. Arp5-si upregulated Brg1-dependent genes to a larger extent compared to Brg1-independent genes (1.77-fold vs. 1.38-fold, *p* = 0.02; Figure 7A). This Brg1 dependency was seen more clearly by comparing the effect of Arp5-si versus Ies6-si and Ino80-si: Arp5-si more effectively increased Brg1-dependent gene expression compared with Ies6-si and Ino80-si, whereas Brg1-independent gene alteration was comparable between them (Figure 7A). Thus, Arp5, but not Ies6 and Ino80, affects the expression of MyoD target genes in a Brg1-dependent manner.

**Figure 7.**
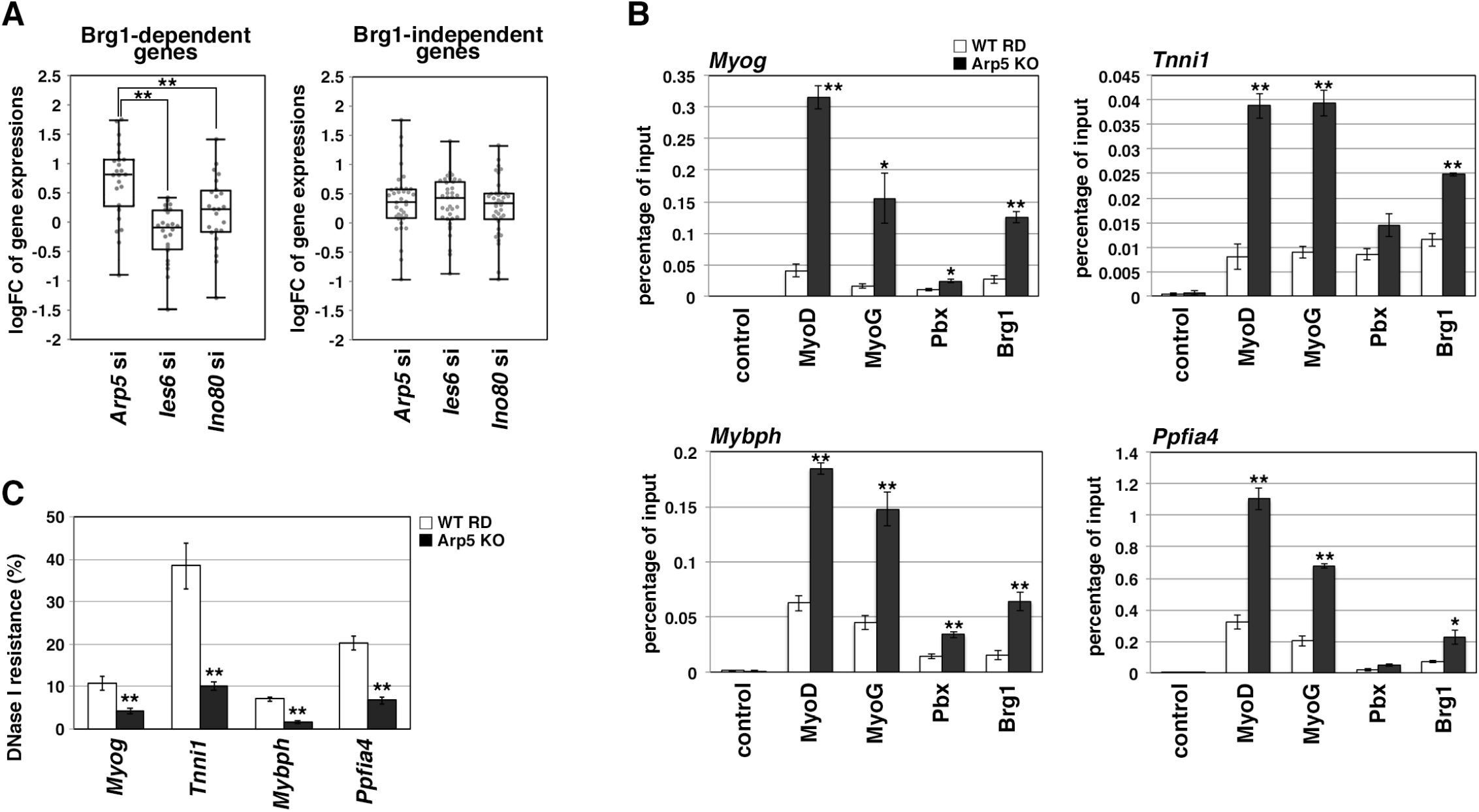
Actin-related protein 5 (Arp5) inhibits the recruitment of MyoD, MyoG, and Brgl-based switch/sucrose nonfermentable (SWI/SNF) to the enhancer region of myogenic genes. (A) Box-and-whisker plot of fold changes (log2) in the expression level of Brgl-dependent and Brgl-independent myogenic genes by Arp5, Ies6, and Ino80 knockdown. (B) Chromatin immunoprecipitation (ChIP) analysis using antibodies against MyoD, MyoG, Pbx, and Brgl in wild-type (WT) and Arp5-knockout (KO) cells. Enrichment efficiency of DNA fragments of MyoG, Tnni1, Mybph, and Ppfia4 enhancer loci was quantified by real-time polymerase chain reaction (PCR). (C) DNase I sensitivity assay of the enhancer loci in WT and Arp5-KO RD cells. All statistical data are presented as the mean± standard error of the mean (SEM). *P < 0.05, **P < 0.01 (Student’s t-test).

To determine the role of Arp5 in regulating Brg1 recruitment to MyoD target loci, we performed chromatin immunoprecipitation (ChIP) assay using Arp5-KO RD cells. MyoD, MyoG, Pbx, and Brg1 were recruited to the proximal promoter region of *Myog*. MyoD, MyoG, and Brg1 significantly accumulated in Arp5-KO cells compared with parental RD cells (Figure 7B). Arp5 KO increased Pbx1 recruitment to a lesser extent. ChIP assay against the enhancer regions of Brg1-dependent myogenic genes, *Tnni1, Mybph*, and *ppfia4*, using public ChIP-Seq and DNase I hypersensitive site (DHS)-Seq databases identified Pbx1- and MyoD-binding regions in their promoter and intronic regions; these regions were DNase I hypersensitive and positive for H3K27Ac, which are markers for the identification of active enhancer regions (Figure 7—figure supplement 1). Pbx–Meis-binding motifs actually exist close to canonical and noncanonical E-box motifs in the focused areas (Figure 7—figure supplement 1). ChIP data showed that all the proteins of interest were recruited to the predicted enhancer regions and that Arp5 KO significantly augmented MyoD, MyoG, and Brg1 accumulation in these regions (Figure 7B).

Finally, we investigated the involvement of Arp5 in chromatin structure alteration of the MyoD/MyoG target loci. DNase I accessibility to the above-mentioned regions was compared between Arp5-KO and parental RD cells (Figure. 7C). The accessibility of all the target regions was significantly higher in Arp5-KO RD cells. The findings illustrate that Arp5 binds to MyoD and MyoG competitively with the Pbx1–Meis1 heterodimer and prevents the recruitment of the Brg1–SWI/SNF complex to their target loci, resulting in chromatin accessibility attenuation and, therefore, transcriptional suppression of myogenic genes.

## Discussion

In this study, we demonstrated a novel role of Arp5 in skeletal muscle differentiation. The H/C region of MyoD is critical for its chromatin remodeling activity. MyoD interacts with some epigenetic regulators, such as SWI/SNF, histone deacetylase (HDAC), p300 histone acetyltransferase (HAT), and p300/CBP-associated factor (PCAF) (Yuan et al., 1996; Puri et al., 1997; Lu et al., 2000; Serna et al., 2001), but how the H/C region contributes to epigenetic regulation is unclear. The Pbx1–Meis1 heterodimer is so far the only identified partner of MyoD-binding directly to this region. Here we reported Arp5 as another binding partner, which competes with the Pbx1– Meis1 heterodimer for binding to the H/C region. The Pbx1–Meis1 heterodimer is believed to recognize the PM motif in the enhancer region of myogenic genes and acts as a pioneering factor by marking this locus for the recruitment of MyoD and its co-factors, such as epigenetic regulators (Cho et al., 2015). Our findings clearly show that the Pbx1–Meis1 heterodimer directly binds to the PM motifs in the promoter region of *MyoG* and *Myf6* and augments the MyoD and MyoG binding to the canonical and noncanonical E-box close to the PM motifs. The ChIP-Seq database and ChIP analysis reveal Pbx1 recruitment to the enhancer regions of some myogenic genes with MyoD. Arp5 inhibits the recruitment of MyoD and MyoG to Pbx1–Meis1-marked loci and, consequently, attenuates the recruitment of chromatin remodelers, such as Brg1– SWI/SNF. Therefore, low Arp5 expression in skeletal muscle tissues contributes to an increase in the chromatin accessibility of MyoD/MyoG target loci and the maintenance of high transcriptional activity of myogenic genes.

Arp5 is a well-known subunit of INO80 and plays an essential role in its ATPase activity and nucleosome sliding (Yao et al., 2015). In HeLa cells knocked down for each subunit of INO80, such as Ino80, Arp8, Ies2, and Ies6, the expression of many genes was altered, and these genes were enriched in several functional pathways such as the p53 signaling pathway, cell cycle, focal adhesion, and extracellular matrix–receptor interaction (Cao et al., 2015). These profiling data are in good agreement with our results using Ino80-si RD cells (Figure 2—figure supplement 1A–E). In osteogenic differentiation of mesenchymal stem cells, INO80 interacts with the WD repeat domain 5 protein and regulates osteogenic marker expression (Zhou et al., 2016). When 11 kinds of INO80 subunits are knocked down in mesenchymal stem cells, all the knockdown cells show a reduction in calcium deposition. Thus, each subunit seems indispensable for the function of INO80. In contrast, we observed many muscle-related genes to be upregulated only in Arp5-si cells but not in Ies6- and Ino80-si cells. Besides, Arp5 alone interrupted MyoD/MyoG–Pbx1–Meis1 complex formation, thereby inhibiting their activities. These results suggest that Arp5 regulates the activities of MyoD and MyoG independently of INO80. We previously demonstrated the INO80-independent role of Arp5 in regulating the phenotypic plasticity of SMCs through direct interaction with myocardin (Morita et al., 2014). Yao et al. (2016) reported that the total abundance of Arp5 in cells is approximately eightfold of that as a component of INO80. Thus, Arp5 seems to have multiple functions besides being a subunit of INO80.

RMS cells are useful for analyzing the antimyogenic function of Arp5 because they highly express Arp5 and MRFs simultaneously. MyoD and MyoG are potent markers of RMS and can induce terminal myogenic differentiation, while their activities are post-transcriptionally dysregulated in RMS (Keller and Guttridge, 2013). Our results provide insight into the post-transcriptional regulation of MyoD and MyoG: abundant Arp5 in RMS induces MyoD and MyoG dysfunction via direct interaction. Arp5 deletion in RD cells leads to a loss of their tumorigenicity in nude mice due to the enhancement of myogenic differentiation. Arp5 is also identified as a T-cell acute lymphocytic leukemia and sarcoma antigen (Lee et al., 2003), although its contribution to carcinogenesis and sarcomagenesis is unclear. In Arp5-si RD cells, the expression of several cancer-associated genes decreased, including those reported to be mutated and abnormally expressed in RMS (Figure 2—figure supplement 1F). Survival data from The Cancer Genome Atlas reveal that high Arp5 expression is unfavorable for the prognosis of liver and renal carcinoma (data not shown). Thus, Arp5 likely has oncogenic properties, in addition to its inhibitory role in myogenesis.

Here we reported that Arp5 expression is remarkably low in both human and mouse skeletal muscle. In contrast, RMS shows increased Arp5 expression with dysregulation of myogenic differentiation compared with tumor-adjacent skeletal muscles. Haldar et al. (2007) reported a synovial sarcoma mouse model in which the ectopic expression of a chimeric SYT-SSX fusion gene was driven from the *Myf5* promoter. In the tumors, Arp5 expression was approximately fivefold upregulated, while several myogenic markers were downregulated. Thus, Arp5 expression seems to be regulated according to the differentiation stage of skeletal muscle lineage cells. Alternative splicing coupled to nonsense-mediated messenger RNA (mRNA) decay (AS-NMD) is important for keeping the expression of Arp5 low in differentiated SMCs (Morita et al., 2014). AS-NMD contributes to post-transcriptional fine-tuning of broad gene expression (Nasif et al., 2018); furthermore, depletion of the splicing factor U2AF35 increased *Arp5* mRNA levels in Hek293 cells (Kralovicova et al., 2015). Tissue-specific alternative splicing is most frequently observed in skeletal muscle, and these splicing events are critical for proper skeletal muscle development and function (Nakka et al., 2018). These facts raise the possibility that AS-NMD contributes to the regulation of Arp5 expression in skeletal muscle and RMS.

This study reported a novel function of Arp5 in myogenic differentiation and tumorigenesis. Arp5 is a novel modulator of MRFs in skeletal muscle differentiation. Our data also cultivated a better understanding of chromatin remodeling events mediated by MRFs and Pbx–Meis in myogenic gene expression. The major limitations of this study are the lack of data on the change in Arp5 expression and its relevance to MRF activation during skeletal muscle development in vivo. Although in vivo experiments were performed using the AAV6 vector, further studies are needed to fully elucidate the mechanism and significance of low Arp5 expression in muscle tissues. It will be also interesting to investigate whether transcriptional and post-transcriptional regulation of Arp5 contributes to physiological and pathological skeletal muscle dysfunction.

## Materials and Methods

### Cell cultures, treatment, and transfections

Human RMS RD cells (supplied by the Japanese Collection of Research Bioresources cell bank), human skeletal myoblasts (Thermo Fisher Scientific), and mouse C3H muscle myoblasts (C2C12 cells; ATCC) were cultured in Dulbecco’s Modified Eagle’s Medium (DMEM) supplemented with 20% FCS. Mouse primary fibroblasts were isolated from the hind limbs of 3-week-old C57BL/6j mice as follows: The extracted hind limb muscle tissues were minced and incubated in a digestion solution (260 U type I collagenase, 3000 PU dispase II, and 10 mM CaCl_2_ in 1 mL of Hanks’ Balanced Salt solution [HBSS]) for 60 min at 37°C. The dispersed cells were washed twice with HBSS and plated on a type I collagen-coated culture dish with DMEM. After 30 min of incubation, the culture medium was removed, and the attached cells were cultured with DMEM supplemented with 20% FCS. The differentiated myoblasts were incubated in differentiation medium (DMEM supplemented with 2% house serum) for 1–3 days.

For myogenic transdifferentiation of the mouse embryo cell line, 10T1/2 cells (ATCC) were treated with 3 µM 5-azacitidine for 24 h and then cultured in DMEM supplemented with 10% FCS for 3–7 days.

To establish Arp5-KO cell lines, RD cells were transfected with the all-in-one Cas9/gRNA plasmid pSpCas9 BB-2A-GFP (PX458; Addgene; gRNA target sequence, ccgttccgcgacgcccgtgccgc). One day after transfection, green fluorescent protein (GFP)-positive cells were sorted and individually cultured. Gene KO in the isolated clones was validated by DNA sequencing and Western blotting.

In knockdown experiments, cells were transfected with predesigned siRNAs using Lipofectamine RNAiMAX (Thermo Fisher Scientific). The siRNA sequences are listed in Table 1. In overexpression experiments, cells were transfected with pCAGGS expression vectors using Lipofectamine 3000 (Thermo Fisher Scientific). The coding sequences of human *ARP5, MYOD1, MYOG, E47, Pbx1a*, and *MEIS1b* were inserted into vectors with FLAG-, HA-, and Myc-tag sequences. For the intramuscular expression of exogenous Arp5, the adeno-associated virus pseudotype 6 (AAV6) expression vector encoding *ARP5* was constructed using AAVpro Helper Free System (AAV6) (TAKARA BIO). Finally, titration of AAV6 particles was performed using the AAVpro titration kit (TAKARA BIO).

**Table 1.**
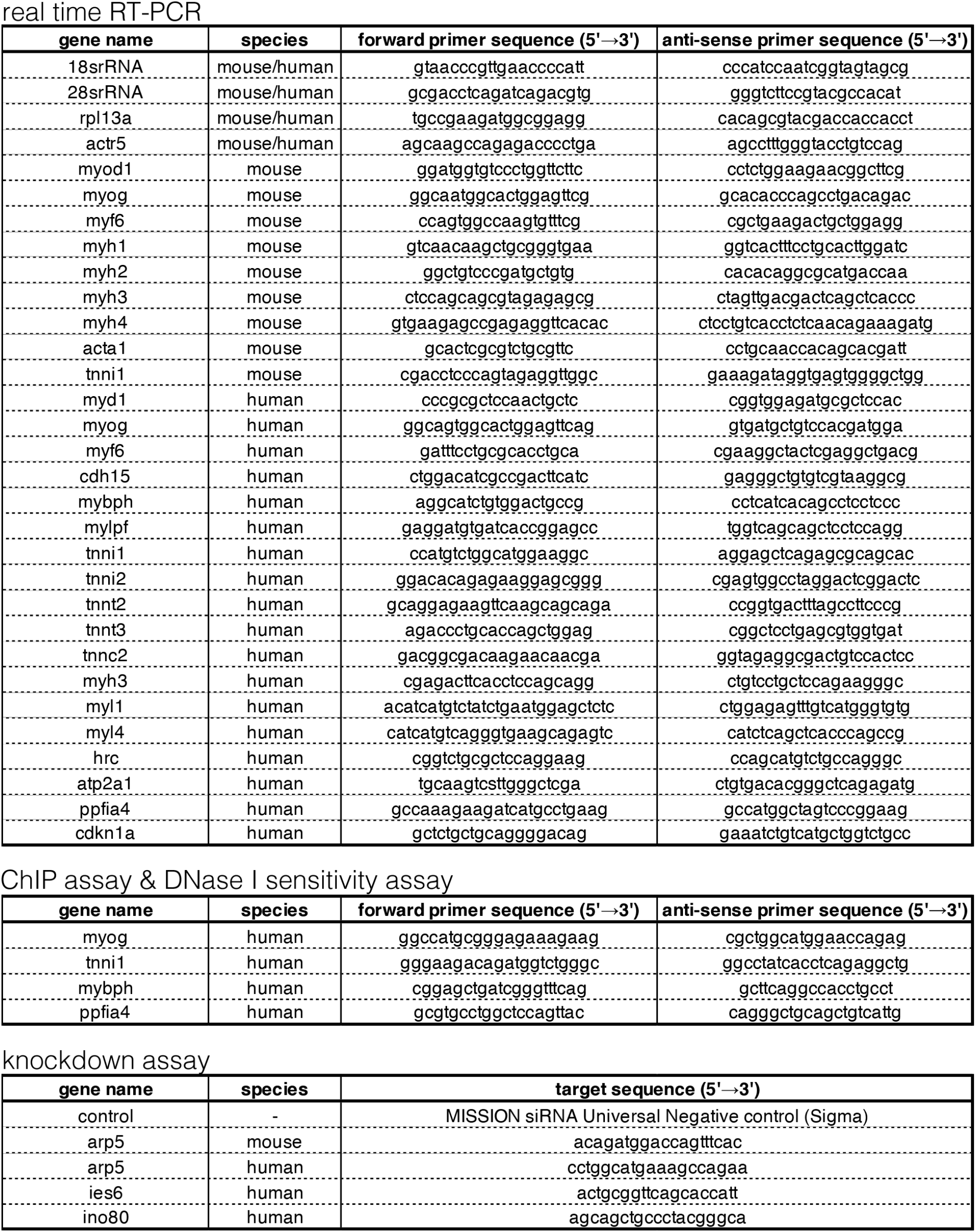
List of PCR primer and siRNA sequences used in this study

### Animal studies

All animal experiments were conducted in accordance with the guidelines for animal experiments specified by the Wakayama Medical University, Japan, and Osaka University School of Medicine, Japan.

For Arp5 overexpression in mouse skeletal muscle tissues, we injected 1 × 10^9^ gv/µL of control-AAV6 (empty vector) or Arp5-AAV6 vector injected into the hind limbs of 5-day-old C57BL/6j mice. Five weeks after injection, the hind limb muscles were extracted, fixed in 10% formalin, embedded in paraffin, and then cut into 5-µm-thick sections. The sections were stained with hematoxylin and eosin and observed.

In xenograft experiments, 2.5 × 10^7^ cells were suspended in 1 mL of an EHS-gel basement membrane matrix (FUJIFILM Wako Pure Chemical Corporation) diluted to 1:1 with phosphate-buffered saline (PBS), and 200 µL of the suspension was subcutaneously inoculated into both sides of the flank of 3-week-old nude mice. The tumor size was measured every week using a pair of calipers, and the tumor volume was estimated as volume = 1/2(length × width^2^). After 6 weeks of inoculation, the mice were sacrificed and xenograft tumors were extracted to obtain total RNA.

### Immunocytochemistry

Cells were cultured on coverslips and fixed with 10% formaldehyde solution. Fixed cells were incubated in blocking solution (0.1% Triton X-100, 0.2% bovine serum albumin [BSA], and 10% normal goat serum in PBS) for 30 min at 37°C and further incubated in a primary antibody solution (1:100 dilution of antibodies in Can Get Signal immunostaining reagent [TOYOBO]) for 2 h. Next, the cells were washed twice with PBS and incubated in a secondary antibody solution (1:400 dilution of Alexa 488– conjugated secondary antibody [Thermo Fisher Scientific] and 1:1000 dilution of Hoechst 33342 [Thermo Fisher Scientific]) in the blocking solution] for 1 h. The cells were again washed twice, mounted on glass slides with Fluoromount (Diagnostic BioSystems), and observed under an all-in-one fluorescence microscope (BZ-9000; Keyence).

### Western blotting

Western blotting was performed, as previously described (Morita et al., 2018). Briefly, total proteins were extracted from cells with 2% sodium dodecyl sulfate (SDS) sample buffer. The proteins were electrophoretically separated using 10% polyacrylamide gels and then transferred to polyvinylidene difluoride membranes. For Western blotting, anti-Arp5 (Proteintech), anti-MyoG (Santa Cruz Biotechnology, Inc.), anti-MyoD (Santa Cruz Biotechnology, Inc.), anti-MYF6 (Santa Cruz Biotechnology, Inc.), anti-MHC (Developmental Studies Hybridoma Bank), anti–glyceraldehyde 3-phosphate dehydrogenase (GAPDH) (Thermo Fisher Scientific), anti-FLAG (Sigma-Aldrich), anti-HA (Roche Applied Science), and anti-Myc (Santa Cruz Biotechnology, Inc.) antibodies were used as primary antibodies.

### Real-time RT-PCR

Total RNA was isolated and then reverse-transcribed using RNAiso Plus (TAKARA BIO) and the PrimeScript RT Reagent Kit with gDNA Eraser (TAKARA BIO). Real-time RT-PCR was performed using the THUNDERBIRD SYBR qPCT Mix (TOYOBO) on the Mic real-time PCR cycler (Bio Molecular System). Nucleotide sequences of the primer sets used in this study are listed in Table 1.

### Reporter promoter analysis

To examine the promoter activity of human *MYOG*, the proximal promoter region (−299/−1) of *MyoG* was isolated by PCR and inserted into the pGL3-basic vector (Promega). The DNA fragments of PME1 and PME2 in the *MyoG* promoter region were also amplified and inserted into the pGL3-γ-actin-TATA vector. These reporter vectors were transfected into cells, together with the pSV-βGal vector (Promega) and the indicated gene expression vectors. After 2 days of transfection, luciferase activity was measured using the Luciferase Assay System (Promega), which was normalized to β-galactosidase activity.

### Co-immunoprecipitation assay

HEK293T cells were transfected with expression vectors to synthesize recombinant proteins. The next day, cells were lysed with 0.5% NP-40, 10% glycerol, and protease inhibitor cocktail (Nacalai Tesque) in PBS and gently sonicated. The lysate was centrifuged to remove cell debris and then incubated with Protein G Sepharose Fast Flow (Sigma-Aldrich) for 1 h at 4°C to remove nonspecifically bound proteins. After centrifugation, the cell lysate proteins were mixed in the indicated combinations and incubated with the ANTI-FLAG M2 Affinity Gel (Sigma-Aldrich) or the Anti-HA Affinity Matrix from rat immunoglobulin G (IgG)_1_ (Sigma-Aldrich) for 2 h at 4°C. The beads were washed thrice with 0.5% NP-40 in PBS, and binding proteins were eluted using SDS sample buffer.

### Protein–DNA pull-down assay

DNA probes for the protein–DNA pull-down assay were generated by PCR using 5’-biotinylated primers and then conjugated to Dynabeads M-280 Streptavidin (Thermo Fisher Scientific). The recombinant proteins synthesized in HEK293T cells were extracted and incubated with the DNA-probe-conjugated beads in 0.5% NP-40, 10% glycerol, and protease inhibitor cocktail (Nacalai Tesque) in PBS for 2 h at 4°C. The beads were washed thrice with 0.5% NP-40 in PBS, and binding proteins were eluted using SDS sample buffer.

### ChIP assay

ChIP assays were performed in WT and Arp5-KO RD cells using the SimpleChIP Plus Enzymatic Chromatin IP Kit (Cell Signaling Technology) according to the manufacturer’s instructions. Anti-MyoD, anti-MyoG, anti-Pbx1/2/3/4, and anti-Brg1 antibodies (all from Santa Cruz Biotechnology, Inc.) were used for immunoprecipitation.

### DNase I sensitivity assay

DNase I sensitivity assays were performed, as previously reported with slight modifications (Gerber et al., 1997). Briefly, WT and Arp5-KO RD cells were suspended in 0.5% NP-40, 10 mM NaCl, 5 mM MgCl_2_, and Tris-HCl at pH 7.4 for 10 min at 4°C to isolate the nuclei. After centrifugation, the pelleted nuclei were resuspended in DNase I buffer (10 mM NaCl, 6 mM MgCl_2_, 1 mM CaCl_2_, and 40 mM Tris-HCl at pH 8.0) and then treated with 1.5 U/50 mL of DNase I (Thermo Fisher Scientific) for 10 min at 37°C. The reaction was stopped by adding an equal volume of 40 mM ethylenediaminetetraacetic acid (EDTA). The digested nuclei were collected by centrifugation, suspended in 50 mM NaOH, and heated for 10 min at 100°C to extract DNA. The resulting solution was diluted 100 times with TE (1 mM EDTA, 10 mM Tris-HCl at pH 8.0). The digestion efficiency of the focused enhancer loci of myogenic genes was calculated by comparing the amount of intact enhancer fragments between DNase-I-treated and DNase-I-untreated samples using real-time PCR.

### DNA microarray

Total RNAs were isolated from RD cells transfected with control, Arp5, Ies6, and Ino80 siRNA for 2 days, using NucleoSpin RNA Plus (TAKARA BIO). mRNAs were reverse-transcribed, and Cy3-labeled complementary RNAs (cRNAs) were synthesized using the Low Input Quick Amp Labeling Kit (Agilent Technologies). The cRNAs were hybridized on a SurePrint G3 Mouse Gene Expression 8 × 60K Microarray (Agilent Technologies), and fluorescence signals were detected using the SureScan Microarray Scanner (Agilent Technologies). The fluorescence intensity was quantified using Feature Extraction Software (Agilent Technologies).

### Statistics and reproducibility

All statistical data were generated from experiments independently repeated at least three times, and values were expressed as the mean ± standard error of the mean (SEM). Data were assessed using Student’s *t*-test, and \**P* < 0.05 and \*\**P* < 0.01 were considered statistically significant.

## Data statement

DNA microarray data have been deposited in the Gene Expression Omnibus (GEO) database (accession no. GSE169681). The data of Arp5 expression profiles in Figure 2A were obtained from the GEO dataset GSE28511.

## Competing Interest Statement

The authors declare no conflicts of interest associated with this manuscript.

## Acknowledgments

We thank all the staff in the Laboratory Animal Center, Wakayama Medical University and the Center for Medical Research and Education of Osaka University Graduate School of Medicine for their technical assistance. We also thank Enago (www.enago.jp) for the English language review. This work was supported by JSPS KAKENHI Grant Number 15K07076 (to T. Morita), 18K06913 (to T. Morita) and 19K07351 (to K. Hayashi).

---

**Figure 1—figure supplement 1.**
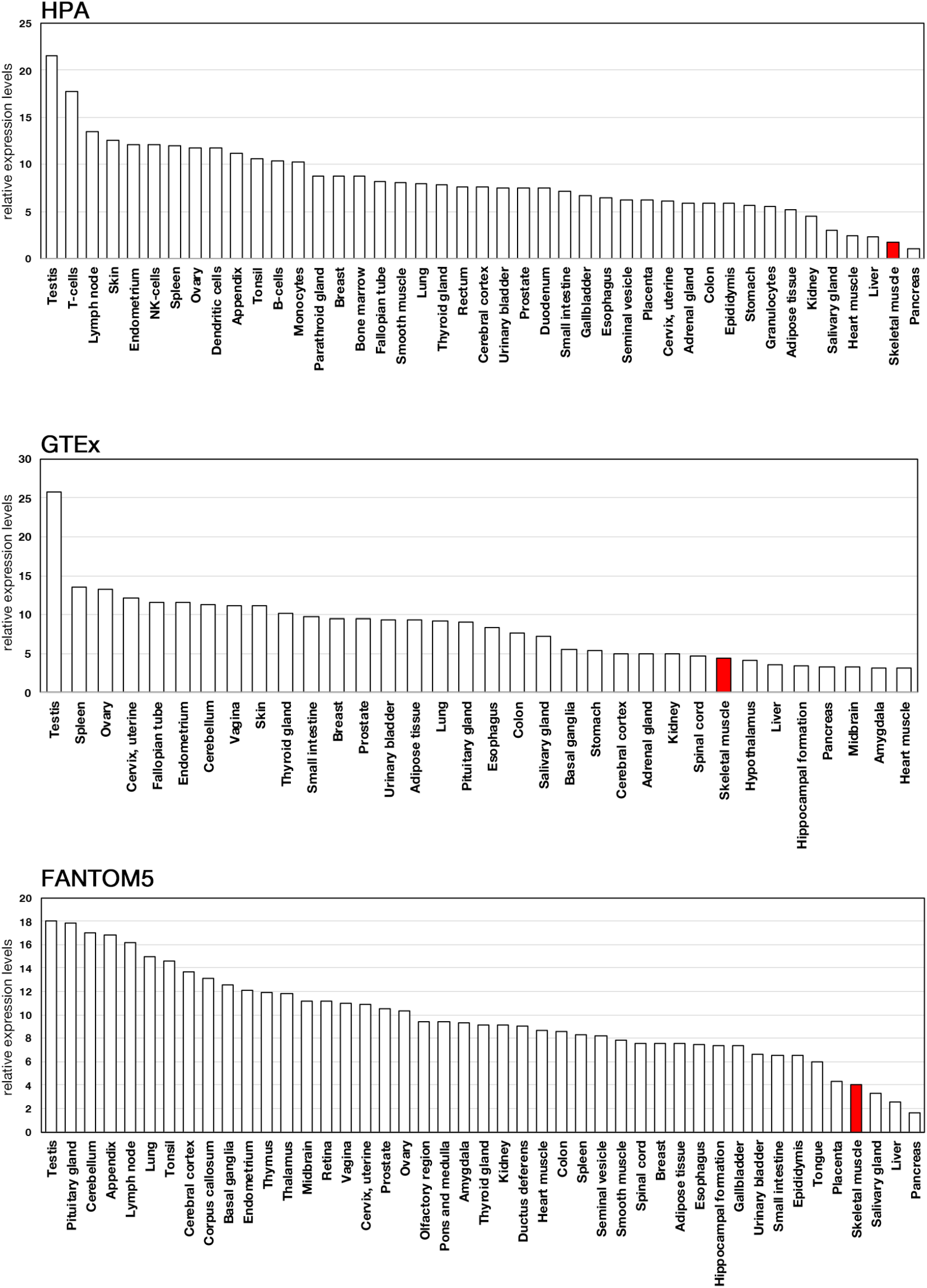
Expression profiles of Actin-related protein 5 (ARP5) in human tissues. Expression data of ARP5 were acquired from three public databases: Human Protein Atlas (HPA) Genotype-Tissue Expression (GTEx), and Functional Annotation of the Mouse/Mammalian Genome 5 (FANTOM5)

**Figure 2—Table supplement 1.** List of the myogenic genes whose expression level is upregulated by Arp5 knockdown in RD cells.

| gene name | relative expression level (vs control siRNA) |  |  |
| --- | --- | --- | --- |
|  | Arp5 siRNA | les siRNA | Ino80 siRNA |
| ACTN3 | 2.076003774 | 0.687842617 | 0.938970893 |
| ATP2A1 | 2.212933663 | 0.64311357 | 0.418973526 |
| CDH15 | 1.835053382 | 1.238002233 | 1.384500699 |
| CKM | 1.747237254 | 0.743827933 | 0.996854717 |
| MEF2B | 1.519753751 | 1.037254304 | 1.03429799 |
| MYBPH | 3.366623852 | 1.222955635 | 1.993441588 |
| MYH7B | 4.734458903 | 0.719439084 | 0.806704257 |
| MYH8 | 2.644230776 | 1.950191916 | 0.185822279 |
| MYLPF | 2.279431645 | 0.962704219 | 1.355377816 |
| MYOG | 1.743530176 | 0.955793102 | 0.734200156 |
| TNNI2 | 3.297109791 | 1.233338111 | 1.630513745 |
| TNNT2 | 2.811110999 | 0.606930828 | 1.855443232 |
| TNNT3 | 2.376308547 | 1.227166077 | 1.549523985 |

**Figure 2—figure supplement 1.**
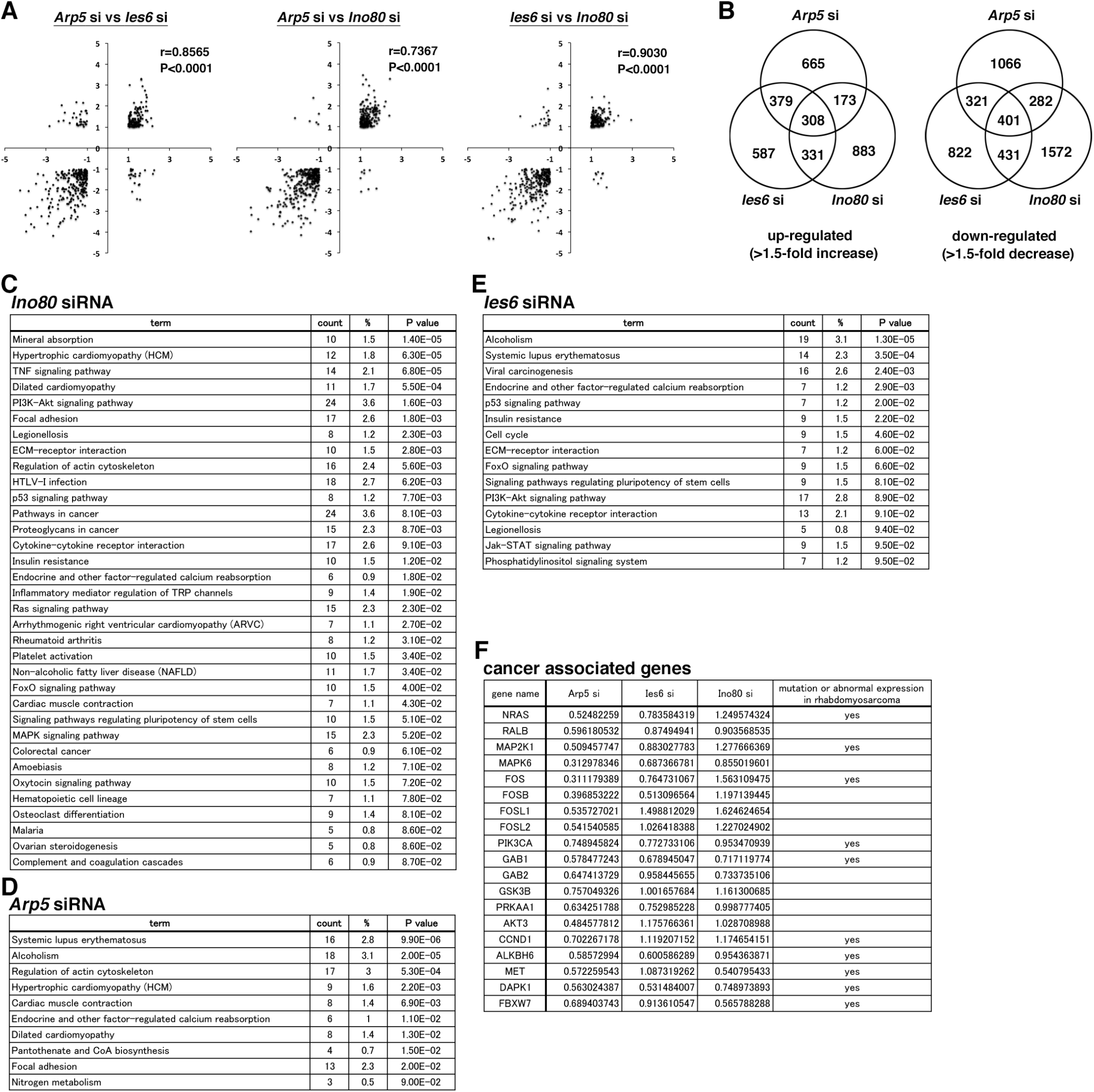
Expression profiles ofgenes altered by Actin-related protein 5 (Arp5), Ies6, and Ino80 knockdown. (A) Scatter plot of fold changes (log2) in the expression level of genes altered by Arp5, Ies6, and Ino80 knockdown. The dataset was filtered for genes with more than twofold increase or decrease. The Pearson’s correlation coefficient (r) was calculated. (B) Venn diagrams of the number of genes whose expression level increased (left) or decreased (right) by more than 1.5-fold by Arp5, Ies6, and Ino80 knockdown. (C) Enrichment analysis of the Kyoto Encyclopedia of Genes and Genomes (KEGG) pathway from the DNA microarray data on Arp5 knockdown in RD cells. (D) Enrichment analysis of the KEGG pathway from the DNA microarray data on Ies6 knockdown in RD cells. (E) Enrichment analysis of the KEGG pathway from the DNA microarray data on Ino80 knockdown in RD cells. (F) List of cancer-associated genes whose expression level was downregulated by Arp5 knockdown in RD cells.

**Figure 4— figure supplement 1.**
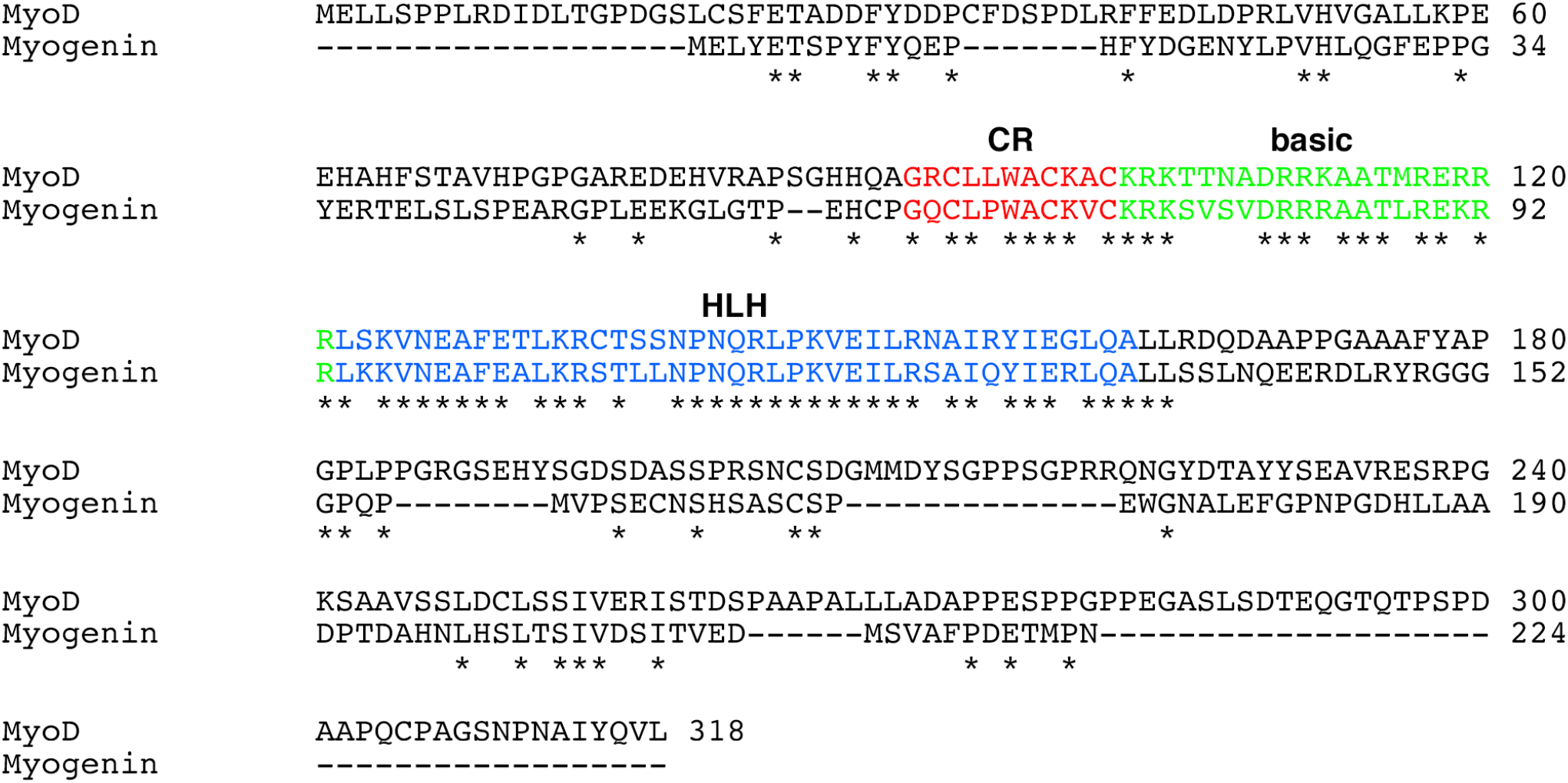
Sequence alignment between human MyoD and MyoG proteins. The CR (red), basic (green), and helix loop helix (HLH) regions are highlighted. Asterisks indicate conserved amino acid residues between them.

**Figure 7— figure supplement 1.**
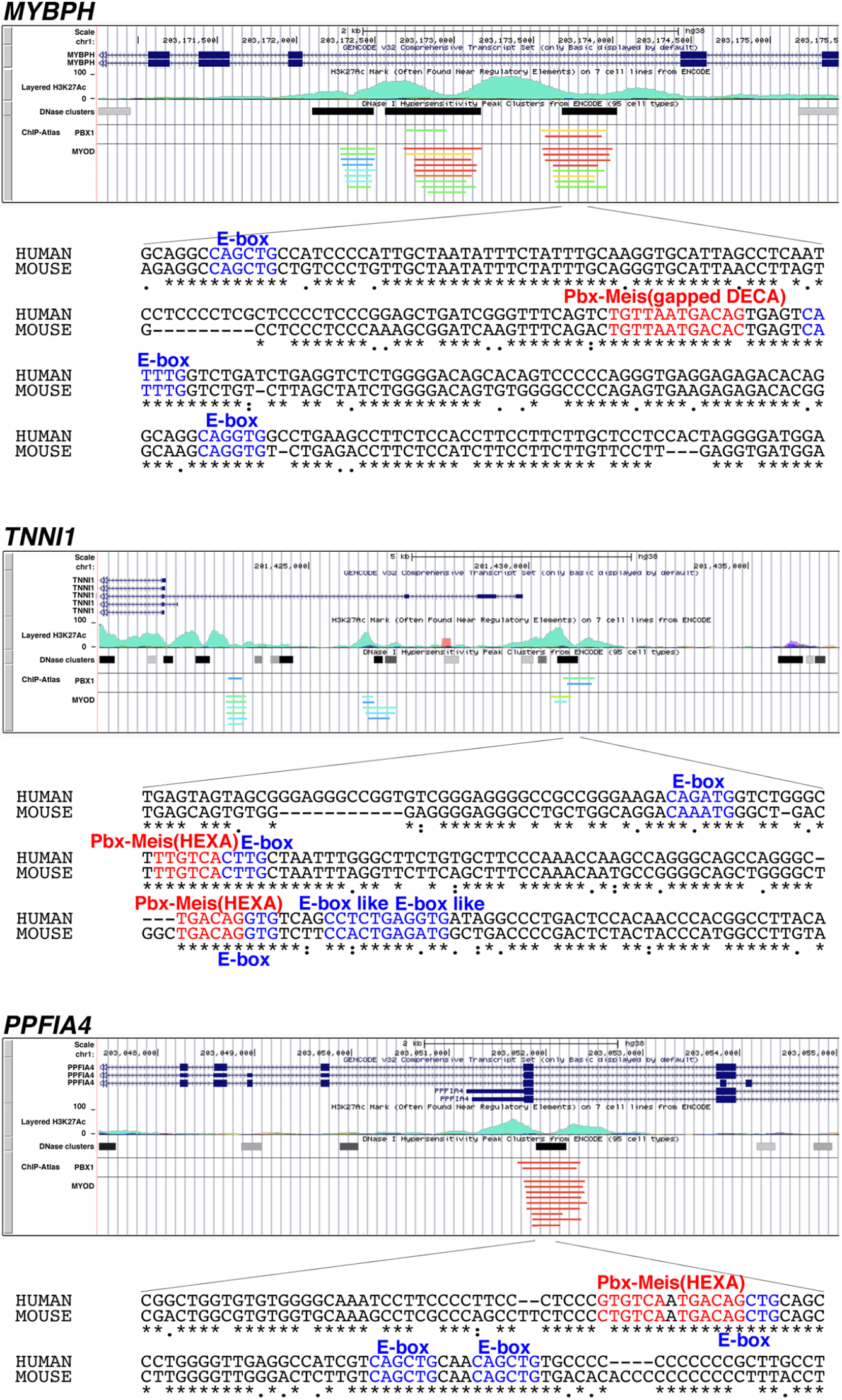
Putative enhancer regions recognized by Pbxl and MyoD/MyoG in human MYBPH, TNNil, and PPFIA4. Data on the enrichment of H3K27 Ac histone markers and DNase I hypersensitivity clusters were acquired from the Encyclopedia of DNA Elements (ENCODE) public database, version 3 (https://www.encodeproject.org). Data on the chromatin immunoprecipitation (ChIP) sequence were acquired from the ChIP-Atlas public database (https://chip-atlas.org). Nucleotide sequences of putative enhancer regions are presented with highlighted Pbx Meis-binding motif (red) and E-box motif (blue). Decameric (DECA, TGAT TGACAG) and hexameric (HEXA, TGACAG) motifs are reported as a consensus sequence of the Pbx Meis-binding site.

## Notes

### Competing Interest Statement

The authors have declared no competing interest.

